# Sex-specific Alterations in Hepatic Cholesterol Metabolism in Young Uteroplacental Insufficiency-induced Low Birth Weight Adult Guinea Pig Offspring

**DOI:** 10.1101/2020.11.12.379891

**Authors:** Ousseynou Sarr, Katherine E. Mathers, Christina Vanderboor, Aditya Devgan, Daniel B. Hardy, Lin Zhao, Timothy R.H. Regnault

## Abstract

**Background:** Intrauterine growth restriction (IUGR) and low birth weight (LBW) have been widely reported as an independent risk factor for hypercholesterolemia and increased hepatic cholesterol underlying liver dysfunction in adulthood. However, the specific impact of uteroplacental insufficiency (UPI), a leading cause of LBW in developed world, on hepatic cholesterol metabolism in later life, is ill defined and is clinically relevant in understanding later life liver metabolic health trajectories.

**Methods:** Hepatic cholesterol metabolism pathways were studied in uterine artery ablation-induced LBW and normal birth weight (NBW) male and female guinea pig offspring at postnatal day 150.

**Results:** Hepatic free and total cholesterol were increased in LBW *versus* NBW males. Transcriptome analysis of LBW *versus* NBW livers revealed that “Cholesterol metabolism” was an enriched pathway in LBW males but not females. Microsomal triglyceride transfer protein and cytochrome P450 7A1 protein, involved in hepatic cholesterol efflux and catabolism, respectively, and catalase activity were decreased in LBW male livers. Superoxide dismutase activity was reduced in LBW males but increased in LBW females.

**Conclusions:** UPI environment is associated with a later life programed hepatic cholesterol accumulation via impaired cholesterol elimination, in a sex-specific manner. These programmed alterations could underlie later life cholesterol-induced hepatic lipotoxicity in LBW male offspring.

**Impact Statement:**

- Low birth weight (LBW) is a risk factor for adult hypercholesterolemia and increased hepatic cholesterol.
- Uteroplacental insufficiency (UPI) resulting in LBW increased hepatic cholesterol content, altered hepatic expression of cholesterol metabolism-related genes in young adult guinea pigs.
- UPI-induced LBW was also associated with markers of a compromised hepatic cholesterol elimination process and failing antioxidant system in young adult guinea pigs.
- These changes, at the current age studied, were sex-specific, only being observed in LBW males and not LBW females.
- These programmed alterations could lead to further hepatic damage and greater predisposition to liver diseases in UPI-induced LBW male offspring as they age.

## Introduction

Cholesterol, is a critical biological molecule, acting as a precursor for the synthesis of steroid hormones, bile acids, and vitamin D, and also being critical as a central modulator of cell membrane proteins, receptor trafficking, signal transduction, and cell membrane fluidity (1). In mammals, the liver is the central organ regulating cholesterol homeostasis through it actions in cholesterol uptake, export, conversion into bile acids, biosynthesis and storage (2–7). When cholesterol homeostasis is disrupted, large concentrations of cholesterol accumulate in the liver resulting in a lipotoxic state associated with oxidative stress and can culminate in nonalcoholic steatohepatitis (NASH) (8–14).

An increase in the consumption of dietary cholesterol and/or genetic susceptibility to hypercholesterolemia can underlie impaired cholesterol homeostasis, cholesterol accumulation and a lipotoxic state in the liver (10,15,16). However, it is now becoming apparent that insults early in the life cycle, such as in prenatal life can also contribute, in a sex-specific manner, to aberrant cholesterol metabolism and hepatic cholesterol overload in adulthood. Animal studies have indicated that a wide variety of experimental *in utero* insults/stress situations (maternal protein undernutrition, dietary restriction, hypoxia, or prenatal nicotine), resulting in IUGR/LBW lead to increased serum and/or hepatic cholesterol in weaned and adult male rat offspring (17–20). Transcriptome analysis in the liver of prenatal maternal food restricted-adult rat offspring revealed that activated cholesterol biosynthesis and downregulated bile acid biosynthesis pathways are main targets for intrauterine programming (21). Studies have highlighted that different models yield different outcomes and therefore, understanding *in utero* insult specific mechanisms is key to identifying both conserved pathways and model-specific alterations which increase our understanding of how these permanent modifications in organ systems and phenotype manifest (22,23). Collectively, this allows for better informed target interventions.

Observational human studies have reported that impaired in-utero growth and the resulting LBW are associated with differentially increased serum concentrations of total and low density lipoprotein (LDL) cholesterol in men and women in later life with sex difference by age groups in adults (24–29). Although, UPI is the leading cause of IUGR/LBW in the developed world (30,31), investigations attempting to understand the specific impact of UPI-induced IUGR/LBW on later life hepatic cholesterol metabolism and cholesterolemia are currently limited to rat models (32,33). New reports have begun to examine the molecular pathways that may underly the association between abnormal fetal growth *in utero* and later life cholesterol metabolism dysregulation in humans, but to date they limited to the analysis of cord blood plasma of IUGR at birth (34), with no follow up on hepatic cholesterol metabolism and associated pathways.

In the present study, we sought to unravel the hepatic metabolic pathways underlying the specific impact of UPI on dysregulated hepatic cholesterol metabolism in young adult female and male LBW offspring. We have chosen to use the well-established pre-clinical animal model of UPI-induced IUGR/LBW in guinea pigs (35), especially given the close similarities between humans and guinea pigs with respect to hepatic and whole-body trafficking and processing of cholesterol (36). With the use of transcriptome analysis, we postulated that LBW offspring, in a sex-specific manner, would display increased hepatic cholesterol by young adulthood in association with alterations in the key regulators of cholesterol metabolism.

## Methods

### Animals

All animal procedures were conducted in accordance with guidelines and standards of the Canadian Council on Animal Care. Animal Use Protocol (AUP-#2009-229) was approved and post approval monitoring conducted by the Western University Animal Care Committee. All investigators understood and followed the ethical principles outlined by Grundy (37), and study design was informed by ARRIVE guidelines (38).

Time-mated pregnant Dunkin-Hartley guinea pigs (Charles River Laboratories, Wilmington, MA, USA) were housed in 12 h light and dark cycles in a temperature (20 ± 2°C) and humidity (30–40%) controlled environment, with access to guinea pig chow (LabDiet diet 5025) and tap water *ad libitum*. All pregnant guinea pigs underwent uterine artery ablation (35,39) at mid-gestation (~32 days, term ~69 days) and sows delivered spontaneously. At the end of the experimental pupping period, male and female pups were defined as normal birth weight (NBW) or low birth (LBW) as per previously reported criteria (39,40). At weaning (postnatal day 15), pups were housed individually in clear perplex containers, with open housing covers and wood chew blocks, on racks allowing sound, smell and visual contact of other animals. From weaning to young adulthood (postnatal day 150), pups were fed *ad libitum* a normal diet (TD.110239; Harlan Laboratories) containing 21.6% kcal protein, 60% kcal carbohydrates and 18% kcal fat. Animal weekly weights were recorded throughout the experiment. At postnatal day (PND) 150, NBW (n = 8) and LBW (n = 8) offspring of each sex were fasted overnight, euthanized via CO2 inhalation (41) in the morning (~ 10 am) of the following day and tissue collection conducted. The whole liver was removed and weighed immediately and the right lobe liver was frozen in liquid nitrogen and stored at −80 °C for later biochemical and molecular analyses.

### Hepatic biochemical analysis

Hepatic triglycerides, total cholesterol, free cholesterol and cholesteryl ester contents were determined with colorimetric assays. Hepatic triglycerides were measured using 50 mg of frozen liver and following the Triglyceride Colorimetric Assay Kit (Item No. 10010303, Cayman Chemical, Ann Arbor, Michigan, USA). Liver content of total cholesterol, free cholesterol and cholesteryl esters was measured as previously described (42,43). Enzymatic reagents for total cholesterol (WAKO Diagnostics: Cholesterol E (CHOD-DAOS method) #439-17501) and free cholesterol (WAKO Diagnostics: Free cholesterol (COD-DAOS) method #435-35801) were used as per manufacturer’s instructions. Cholesteryl ester was determined as the difference between total cholesterol and free cholesterol.

### RNA isolation

Total RNA for microarray analysis was isolated from a sub cohort of five NBW and five LBW livers of each sex at the Genome Québec Innovation Centre (Montreal, QC) while independent validation of microarray data was performed using RNA extracted from all NBW and LBW animals of the experiment, including the microarray cohort. For the microarray analysis, the frozen liver was pulverised using a mortar and pestle then 50 mg homogenized in 1 mL of Trizol (Invitrogen, Burlington, ON, Canada). Following a 5 min incubation at room temperature, the homogenate was treated with 100 μL of gDNA eliminator solution then shaken vigorously and maintained at room temperature for 2-3 min. The homogenate was centrifugated at 12,000 g for 15 min at 4 °C and the upper aqueous phase transferred to a clean 2 mL safe-lock microcentrifuge tube. RNA in the aqueous phase was then extracted using the RNeasy Plus Universal Mini Kit (Qiagen, Toronto, ON, Canada) with the Qiacube instrument (Qiagen), following the manufacturer’s instructions. Total RNA for validation of microarray data was isolated from frozen ground liver (100 mg) using Trizol reagent (Invitrogen) and the Purelink RNA Mini Kit according to manufacturer’s instructions (Invitrogen). Total RNA for both microarray analysis and validation of microarray data was quantified using a NanoDrop Spectrophotometer 2000 (NanoDrop Technologies, Inc., Wilmington, DE, USA). RNA integrity was assessed using a 2100 Bioanalyzer (Agilent Technologies, Waldbronn, Germany). Only RNA with RIN > 7 were considered for the analyses.

### RNA labelling, microarray hybridization, scanning and processing

A whole-Transcript Expression Analysis Gene Titan was conducted (Genome Québec Innovation Centre, QU, Canada). Sense-strand cDNA was synthesized from 100 ng of total RNA, and fragmentation and labeling were performed to produce ss-cDNA with the GeneChip^®^ WT Terminal Labeling Kit according to manufacturer’s instructions (Thermo Fisher Scientific, Waltham, MA, USA). After fragmentation and labeling, 2.1 μg cDNA was hybridized onto Guinea Pig Gene 1.1 ST Array Plate (Thermo Fisher Scientific) and processed on the GeneTitan^®^ Instrument (Thermo Fisher Scientific) for Hyb-Wash-Scan automated workflow. The microarray scanned images were imported into the Transcriptome Analysis Console (TAC) (version 4.0) (Life Technologies, Carlsbad, CA, USA) and the raw microarray data (.CEL files) were normalized using the Robust Multiple-Array Averaging (RMA) method which is implemented in the TAC software. The RMA method converts .CEL files to .CHP and summarizes the probes into a single signal for each probeset. Differentially expressed genes were identified using the TAC Expression (Gene) analysis method. Microarray data have been deposited in the National Center for Biotechnology information Gene Expression Omnibus (GEO; accession number GSE161124).

### Quantitative reverse transcription PCR (RT-qPCR)

Independent validation of microarray data was performed by examining levels of selected differentially expressed transcripts using RT-qPCR. cDNA synthesis was performed with 1 μg of total RNA using the MultiScribe™ Reverse Transcriptase Kit (Thermo Fisher Scientific). Primers were designed from *cavia porcellus* sequences available in Ensembl or NCBI databases using the NCBI/Primer-BLAST tool. Detailed information for primer sequences (forward and reverse) is provided in **Supplementary Table 1**. Amplification reactions and disassociation curves of 4-fold dilution series of cDNA were carried out on a CFX384 machine (Bio-Rad, Bio-Rad, Mississauga, ON), for determination of primer efficiencies. All reaction efficiencies were between 90 and 100 % with correlation coefficients of > 0.95. GAPDH and beta actin genes were simultaneously used as internal control genes for normalization. All measurements contained a negative control (no cDNA template). Each RNA sample was analyzed in triplicate. The 2^-ΔΔCt^ method was used to determine the relative expression of each transcript. The mean levels of each target transcript in LBW were divided by mean level in NBW to indicate the fold change for comparisons to array results.

### Immunoblot analysis

Proteins were extracted from frozen ground liver (50 mg) in radioimmunoprecipitation assay (RIPA) buffer as previously described (44). Equal amounts of total proteins (20 μg) were separated on a 7.5 or 10% SDS-polyacrylamide gel and transferred onto PVDF membranes. Membranes were then blocked with 5% non-fat milk in 0.1% TBS-Tween-20 and probed with primary antibodies against Low-density lipoprotein receptor (Ldlr, MABS1208; 1:1000), fatty acid synthase (Fas, sc-20140; 1:1000), Acetyl-CoA carboxylase (Acc, C83B10; 1/1000), Cytochrome P450 Family 7 subfamily A member 1 (Cyp7a1, ab65596; 1:1000), ATP binding cassette subfamily G member 8 (Abcg8, sc-30111; 1:1000), Sterol regulatory element-binding protein 2 (Srebp2, sc-5603; 1:1000), Apolipoprotein E (Apoe, sc-98574; 1/500), Microsomal triglyceride transfer protein (Mtp, ab63467;1:1000), Superoxide dismutase 1 (SOD1, 2770S; 1/500) and Catalase (CAT, sc-69762; 1/500). Membranes were finally incubated with HRP-conjugated anti-rabbit (7074S; 1:10000; Cell Signaling Technology), anti-mouse (7076S; 1:5000, Cell Signaling Technology). Membranes were developed using the Clarity or Clarity Max ECL substrate (BioRad). The chemiluminescence signal was captured with the ChemiDoc MP Imaging System (BioRad), and protein band densitometry was determined using the Image Lab software (Bio-Rad). Ponceau staining was used as the loading control, as previously published (45,46).

### SOD, CAT, GSH and GSSG assays

Superoxide dismutase (SOD; kit n° 706002, Cayman Chemical) and catalase (CAT; kit n° 707002, Cayman Chemical) enzyme activities, reduced glutathione (GSH; kit n° 707002, Cayman Chemical) and oxidized GSH (GSSG; kit n° 707002, Cayman Chemical) in livers were measured following kit instructions. Briefly, 50 mg of ground liver were homogenized in 500 μL of sucrose-mannitol buffer (70 mM sucrose, 210 mM mannitol, 20 mM HEPES and 1mM EGTA) and centrifuged at 1, 500 × g for 5 minutes at 4°C. The SOD activities in supernatants were subsequently measured at 450 nm and cat activities were assayed at 540 nm in diluted supernatants (1:2000). SOD and CAT activities were expressed relative to total protein concentration for each homogenate. For GSH and GSSG measurements, liver tissue (50 mg) was homogenised in 1 mL of 2-(N-morpholino)ethanesulphonic acid (MES) buffer provided in the kit. The homogenate was centrifuged at 10, 000 x g for 15 min at 4°C and the supernatant collected. The supernatant was deproteinized according to the kit technical manual. The deproteinized samples were then diluted 30 times and used for total GSH (GSH+GSSG) measures. For the exclusive measurement of GSSG, deproteinized and diluted (1:5 dilution) samples as well as standards were derivatized with addition of 10 uL of 2-vinylpyridine 1M per mL of sample and 1h incubation at room temperature. Total GSH and GSSG were measured at 405 nm at 5 min intervals for 30 min. GSH standard curves from 0 to 16 μM and 0 to 8 for total gsh and gssg experiment, respectively, were employed. The GSH was calculated by subtracting GSSG from total GSH.

### Statistical analysis

#### Non-microarray data

A two-way ANOVA was used to determine the main effect of birth weight, sex and possible interactions, followed by Bonferroni *post hoc* test using GraphPad 8 (GraphPad Software, San Diego, CA, USA). Student’s t-test was performed when appropriate. Data are presented as mean ± SEM and a probability of 5% (P< 0.05) was considered significant for all analyses.

#### Microarray data

Only genes that met the criteria of |fold change| >= 2 and p-value < 0.05 were defined as differentially expressed. To explore biological processes and pathways associated with the differentially expressed genes, we performed gene ontology (GO) biological process term enrichment (adjusted P < 0.05) using *human* as target organism and the g:GOSt functional profiling in g:Profiler web server (https://biit.cs.ut.ee/gprofiler/gost).

## Results

### Phenotypic traits

By using diathermy partial ablation of branches of the uterine artery, UPI was induced to generate LBW, as described previously (35). Those pups that at birth were < 85 grams, were classified as LBW and as a group weighed less than sex-matched NBW pups (p < 0.0001; **Table 1**). At the time of tissue collection (PND 150; **Table 1**), LBW and NBW offspring had similar mean body and absolute and relative liver weights within each sex group. However, at this age, females displayed overall lower body, absolute and relative liver weights than males, independent of the birth weight (p < 0.05; **Table 1**). Male offspring overall exhibited higher hepatic free cholesterol and total cholesterol than females (p < 0.05; **Table 1**). Interestingly, hepatic free cholesterol and total cholesterol contents was increased in LBW males compared to NBW males (p < 0.05; **Table 1**). No significant difference was observed for hepatic free cholesterol and total cholesterol content in NBW or LBW females (**Table 1**). Birth weight or sex at PND 150 had no impact upon hepatic cholesteryl ester and triglyceride contents (**Table 1**).

**Table 1:**
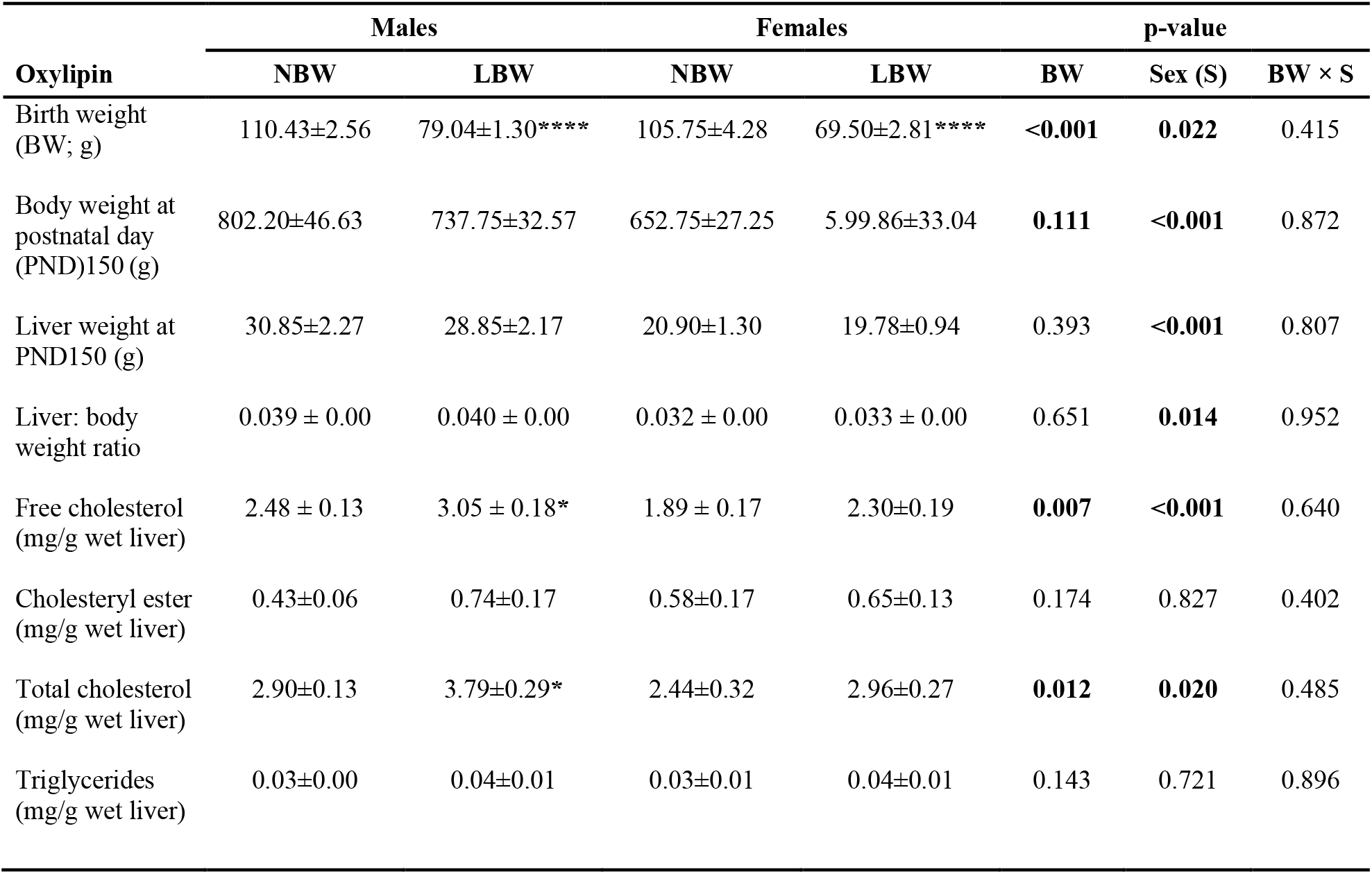
Phenotypic traits of the offspring

| Oxylipin | Males |  | Females |  | p-value |  |  |
| --- | --- | --- | --- | --- | --- | --- | --- |
|  | NBW | LBW | NBW | LBW | BW | Sex (S) | BW × S |
| Birth weight (BW; g) | 110.43±2.56 | 79.04±1.30**** | 105.75±4.28 | 69.50±2.81**** | <0.001 | 0.022 | 0.415 |
| Body weight at postnatal day (PND)150 (g) | 802.20±46.63 | 737.75±32.57 | 652.75±27.25 | 599.86±33.04 | 0.111 | <0.001 | 0.872 |
| Liver weight at PND150 (g) | 30.85±2.27 | 28.85±2.17 | 20.90±1.30 | 19.78±0.94 | 0.393 | <0.001 | 0.807 |
| Liver: body weight ratio | 0.039 ± 0.00 | 0.040 ± 0.00 | 0.032 ± 0.00 | 0.033 ± 0.00 | 0.651 | 0.014 | 0.952 |
| Free cholesterol (mg/g wet liver) | 2.48 ± 0.13 | 3.05 ± 0.18* | 1.89 ± 0.17 | 2.30±0.19 | 0.007 | <0.001 | 0.640 |
| Cholesteryl ester (mg/g wet liver) | 0.43±0.06 | 0.74±0.17 | 0.58±0.17 | 0.65±0.13 | 0.174 | 0.827 | 0.402 |
| Total cholesterol (mg/g wet liver) | 2.90±0.13 | 3.79±0.29* | 2.44±0.32 | 2.96±0.27 | 0.012 | 0.020 | 0.485 |
| Triglycerides (mg/g wet liver) | 0.03±0.00 | 0.04±0.01 | 0.03±0.01 | 0.04±0.01 | 0.143 | 0.721 | 0.896 |

### Hepatic Transcriptome

Analysis of whole hepatic transcriptome results identified that between LBW and NBW offspring livers, 51 (29 up-regulated and 22 down-regulated) and 31 (8 up-regulated and 23 down-regulated) genes were differentially expressed (|fold change| >= 2 and p-value < 0.05) in males and females respectively (**Fig. 1; Supplementary Tables 2** and **3**). The Venn diagram (**Fig. 1A**), the volcano plots (**Fig. 1B, C**) and 2D hierarchical clustering chart/heatmaps (**Fig. 2A, B**) indicate the degree of separation among the LBW and NBW groups in males and females. Only two genes encoding thioredoxin pseudogene and small nucleolar RNA SNORA2/SNORA34 family transcripts were differentially expressed between LBW and NBW in both males and female offspring (**Fig. 1A**).

**Fig. 1:**
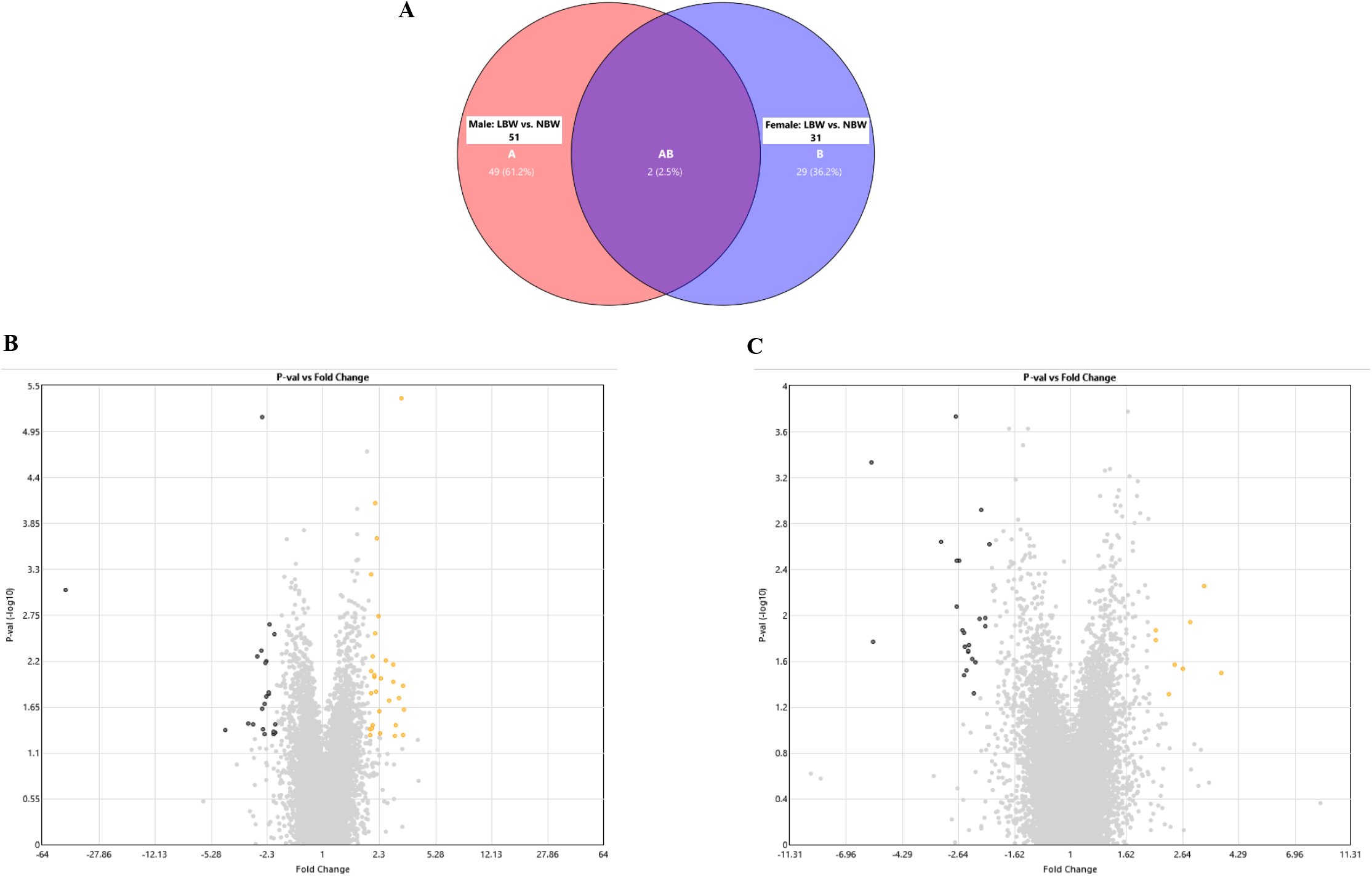
Differential gene analysis, volcano plots and hierarchical clustering chart/heat maps. (**A**) Venn diagram demonstrating differentially up- and down-regulated genes in 5 NBW *versus* 5 LBW offspring using Transcriptome Analysis Console (TAC) Software at a |fold change| >= 2 and p-value < 0.05. Volcano plots were generated to visualize the differentially expressed genes (DEGs) between LBW and NBW in males (**B**) and females (**C**). The x-axis indicates the fold-change and the y-axis represents the log10 p-values. The orange and red points indicate the up-regulated and down-regulated mRNAs with statistical significance, respectively.

**Fig. 2:**
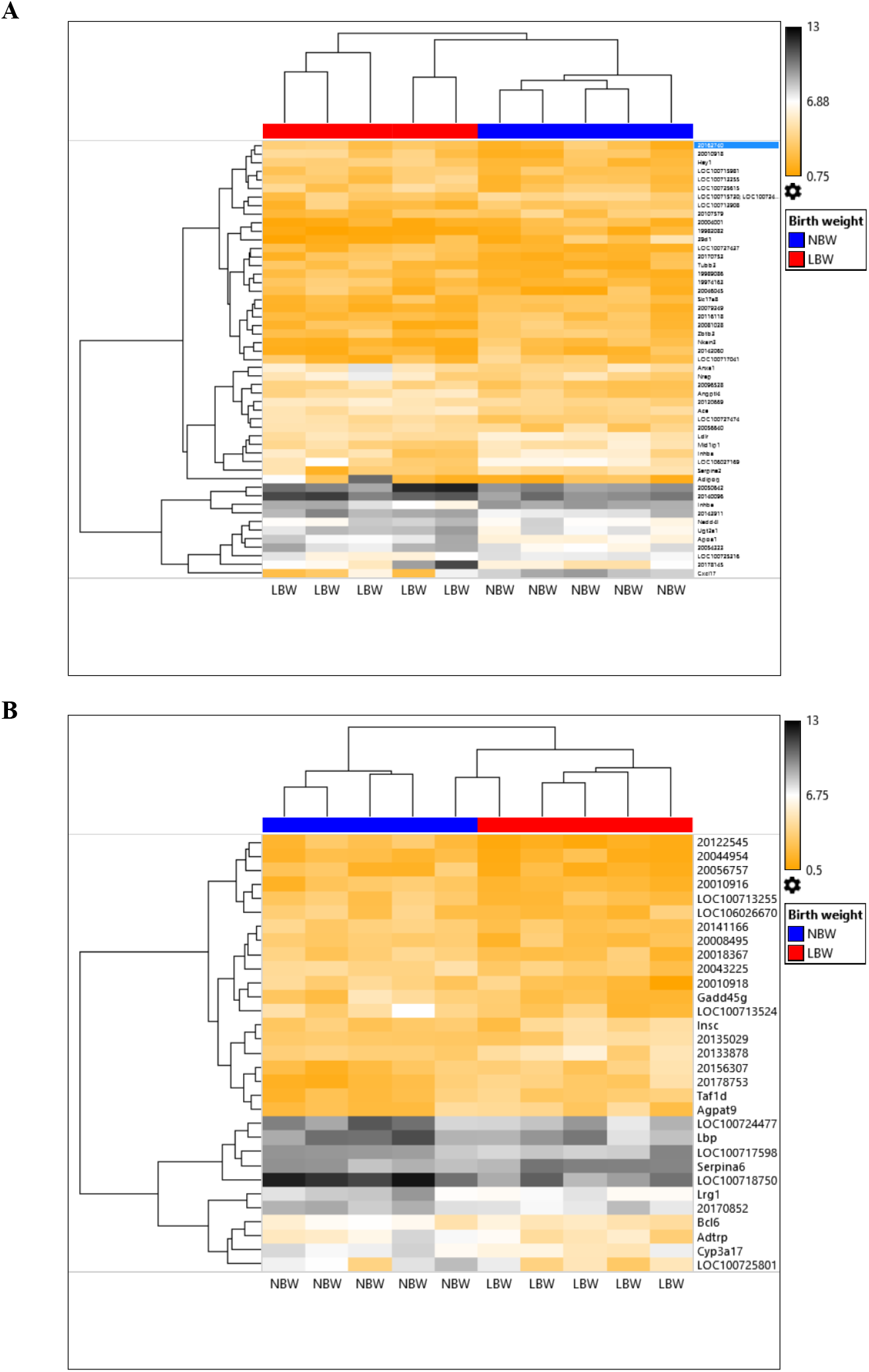
Heat map looking at differential changes of genes. Hierarchical clustered heat maps of DEGs between LBW and NBW in males (**A**) and females (**B**). Each column represents one sample and Value to the right of colour scale indicates the (log2) gene expression. The intensity of the colour indicates the expression level, with black representing a high expression level and orange representing a low expression level. Gene symbols are indicated on the right of the heat maps.

### Functional analysis of the hepatic transcriptomic profile and validation of microarray data

In the biological process analysis of the differentially expressed genes in LBW *versus* NBW males, 18 enriched gene ontology (GO) biological processes were observed, with positive regulation of hepatic fatty acid metabolism process, lipid transport, lipid localization, regulation of fatty acid biosynthetic process and positive regulation of lipid metabolic process being the five top ranked processes (**Fig. 3A, Table 2**). No significant changes to GO biological processes were observed in LBW females (**Fig. 3B**). The KEGG functional enrichment analysis identified that cholesterol metabolism was a significantly enriched pathway in LBW males (adjusted p < 0.05. **Fig. 3A**). Specifically, in LBW males, the reverse cholesterol transport-related genes (*Apoa1* and *Angplt4*) were up-regulated while *Ldlr* gene associated with the internalizing of circulating LDL cholesterol, was down-regulated (**Table 2, Supplementary Table 2**).

**Fig. 3:**
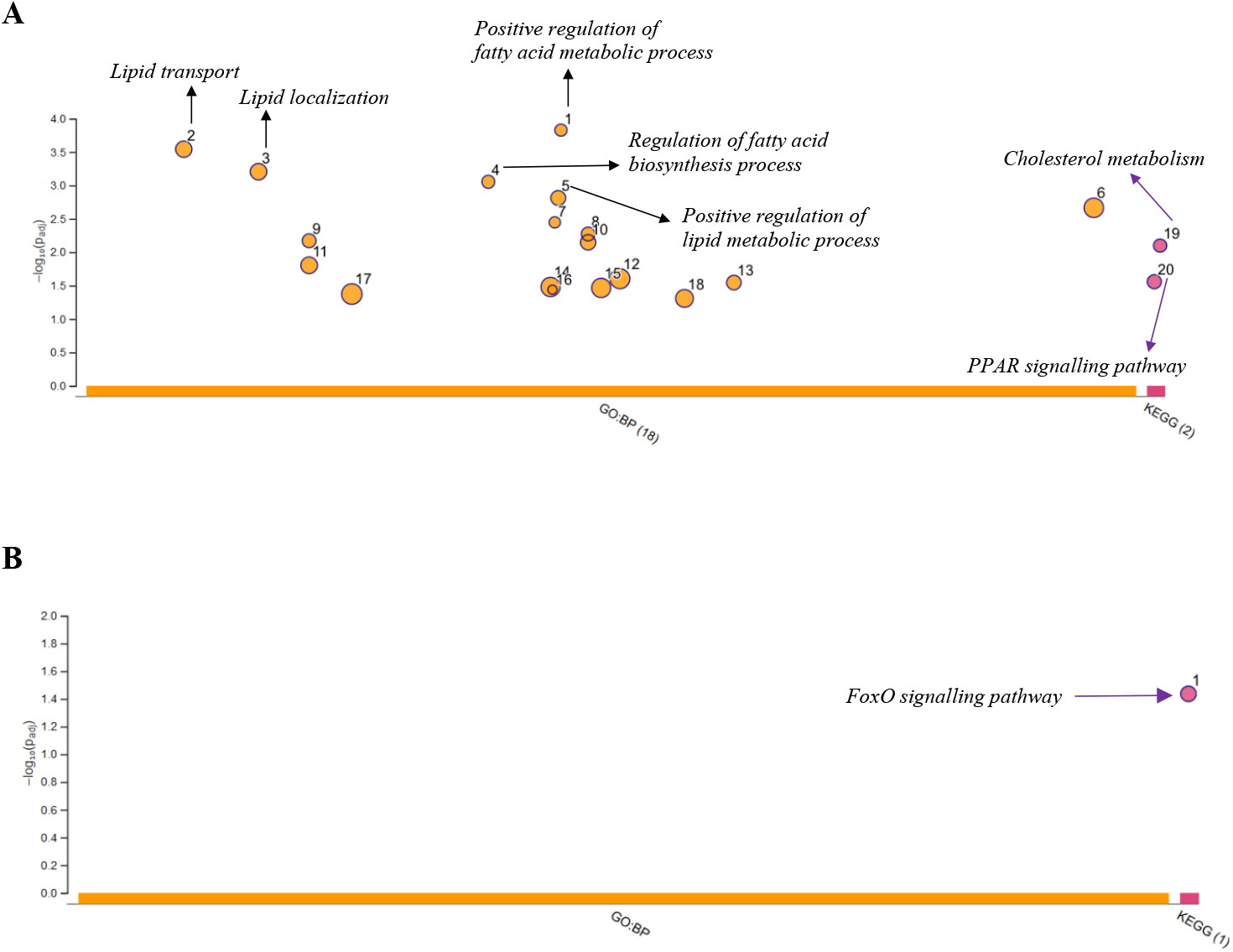
Gene function analysis. Manhattan plots illustrate the enrichment analysis results on DEGs between LBW and NBW in males (**A**) and females (**B**). The x-axis represents functional terms that are grouped and colour-coded by data sources (e.g. biological processes (BP) from gene ontology (GO) are orange circles and enriched KEGG pathways are purple circles). The circle sizes are in accordance with the corresponding term size, i.e. larger terms have larger circles. The y-axis shows the adjusted enrichment p-values in negative log10 scale.

**Table 2.**
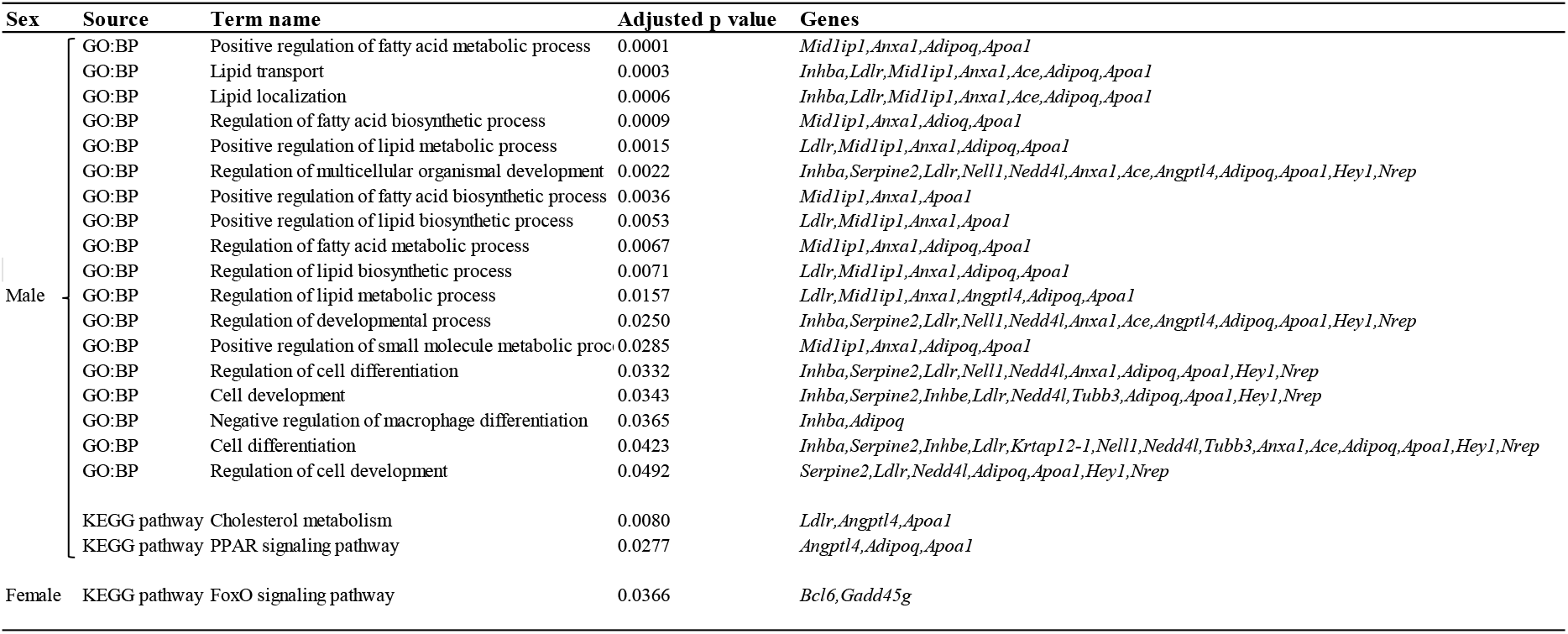
Gene ontology (GO) terms and KEGG pathways overrepresented among differentially expressed genes in livers of LBW versus NBW offspring

The PPAR signalling pathway was also an up-regulated pathway in LBW versus NBW offspring (adjusted p < 0.05. **Fig. 3A**). This latter pathway included *Adipoq* in addition to *Apoa1* and *Angplt4* (**Table 2, Supplementary Table 2**). FoxO signalling pathway was the only significantly enriched pathway for differentially expressed genes in LBW females (adjusted p < 0.05. **Fig. 3B**). This pathway included down-regulated *Bcl6* and *Gadd45g* (**Table 2, Supplementary Table 2**).

Genes involved in cholesterol metabolism, PPAR and FoxO signalling pathways and other differentially expressed genes not related to these pathways were selected to be validated by RT-qPCR independent RNA samples (**Fig. 4**). *Apoa1* gene was increased 4.76-fold (*p* = 0.037) in LBW relative to NBW males, similar to the fold change in the microarray data of 2.92-fold (p = 0.037). *Angplt4* was also confirmed to be increased by 3.7 (*p* = 0.023) similar to microarray fold change of 2.84 and 1.2 (p = 0.011). In LBW versus NBW males, *Ldlr* and *Gstt2* genes were also confirmed to be decreased by −2.16 and −3.05-fold (*p* = 0.008 and 0.020) and similar to microarray fold changes of −2.39 and −2.44, respectively (p = 0.021 and 0.041). *Inhba*, which was −2.82-fold lower in the microarray data (p = 0.036), was relatively lower in the LBW male livers by 2.89-fold (*p* = 0.061). Although not significant, *Anxa1*, which was 2.18-fold higher in the microarray data (p = 0.015), was relatively higher in the LBW male livers by 1.72-fold (*p* = 0.540). *Lbp* was significantly lower by −2.28-fold in the LBW female livers in the validation cohort (*p* = 0.038), similar to the −2.96-fold decreased in the LBW microarray female cohort (p = 0.026). Lastly, while down-regulated in LBW females in the microarray data, *Lcn2* and *Bcl6* tended to decrease in the validation cohort (p = 0.095 and 0.180, respectively).

**Fig. 4:**
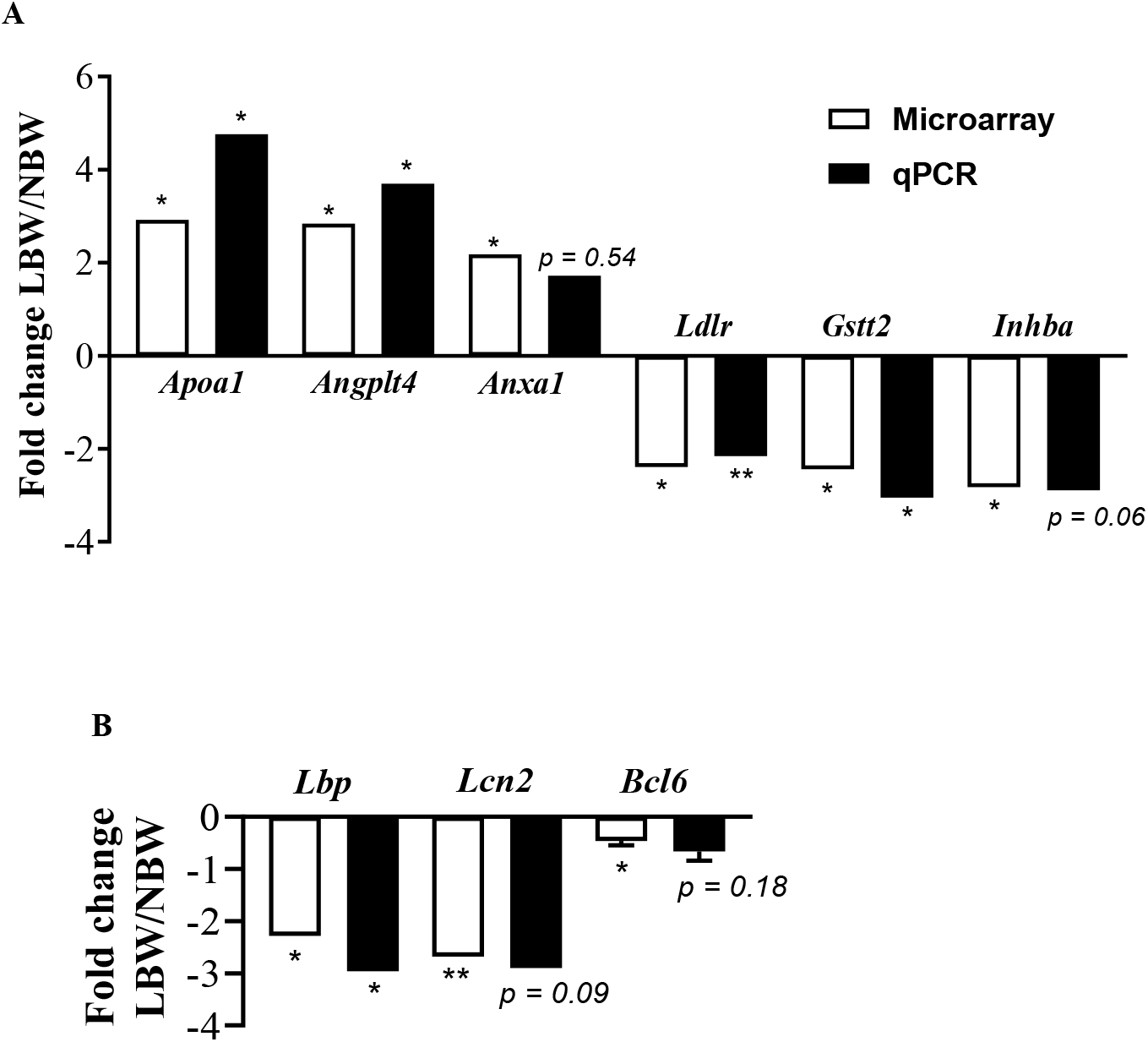
RT-qPCR validation for selected differentially expressed genes. Transcriptomic differences between NBW and LBW in males (**A**) and females (**B**) were evaluated by RT-qPCR. RT-qPCR fold change values are the ratios of LBW to NBW; ratios are inversed and preceded by a minus sign for value less than 1 (i.e., a ratio of 0.5 is expressed as −2). n = 7-8 samples per group. Data are mean ± SEM. *p < 0.05 and **p < 0,01 using Student t-test for independent samples.

### Hepatic content of proteins related to cholesterol metabolism

Selected proteins were studied on *a priori* based on their known biological role in hepatic cholesterol metabolism. The levels of hepatic proteins involved cholesterol uptake (Ldlr, Apoe), biosynthesis (Srebp2), catabolism (Cyp7a1), efflux into bile acids (Abcg8) and export to blood (Fas, Acc, Abca1 and Mtp) were determined in LBW and NBW livers (**Fig. 5**). Female livers displayed significantly higher levels of Ldlr and Srebp2 protein than males (p < 0.01). Neither birth weight nor sex impacted Apoe protein. Females displayed lower Mtp and Acc protein levels than males (p <0.01) and Fas protein levels were not affected by birth weight nor sex. Cyp7a1 and Mtp protein levels were both reduced in male LBW compared to male NBW livers.

**Fig. 5:**
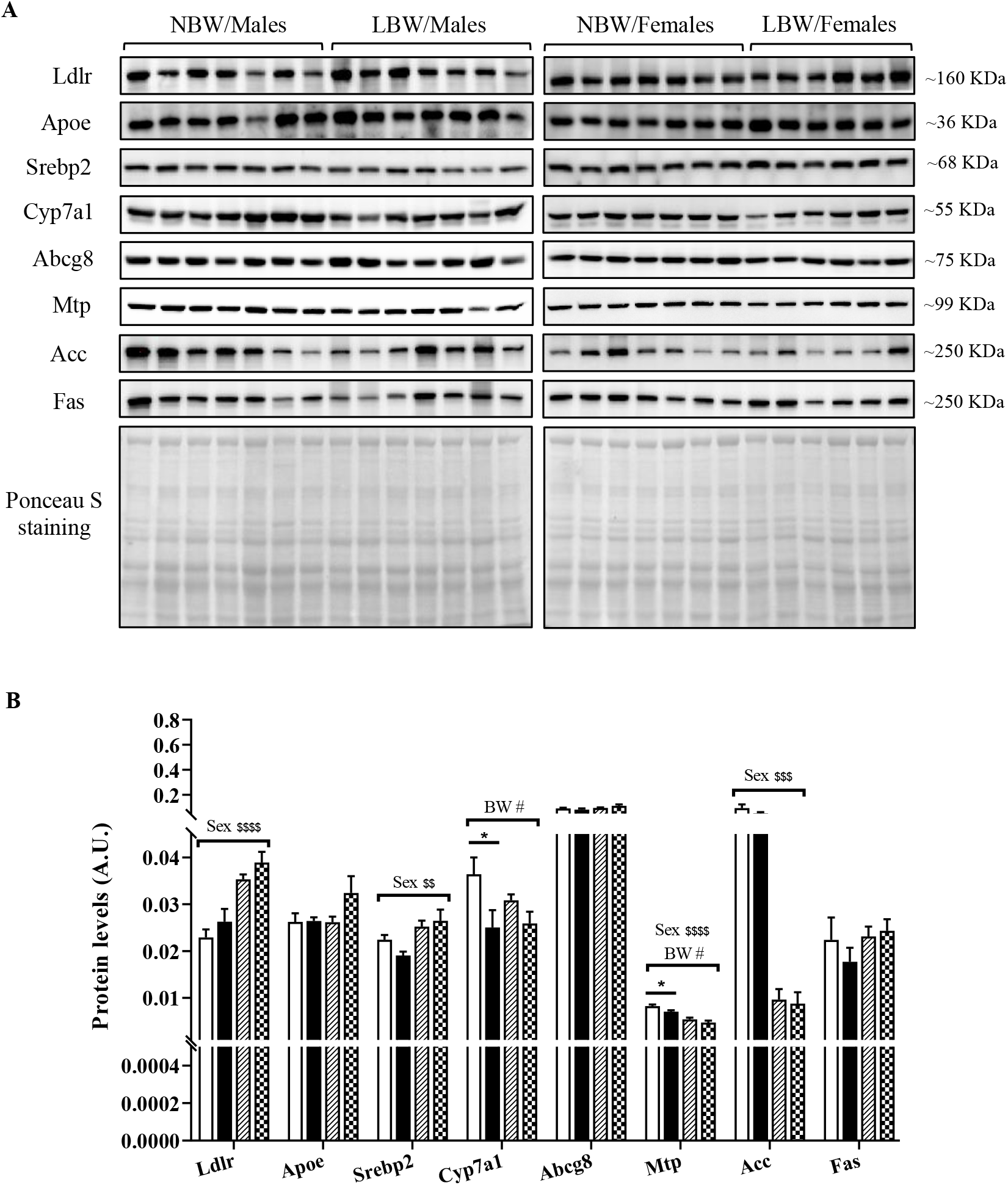
Immunoblotting for proteins related to cholesterol metabolism. Panel **A** shows representative blots of the targeted proteins and ponceau S as a loading control. Panel **B** indicates normalized densitometry values (targeted protein: ponceau) of targeted proteins. Data are mean ± SEM of 6 to 7 animals per birth weight/sex group. ^$$^p <0.01 and ^$$$$^p < 0.0001 for the main effect of sex, ^#^p <0.05 for the main effect of birth weight, using a two-way ANOVA. *p < 0.05 when comparing NBW/Males *vs*. LBW/Males by Bonferroni *post hoc* test.

### Hepatic antioxidant systems

As elevated hepatic cholesterol is associated with oxidative stress (9,10), we quantified markers of this latter. The protein levels and activities of antioxidant enzymes SOD and CAT, as well as GSH and GSSG concentrations were determined in LBW and NBW livers. Sex had a significant effect on SOD1 protein level, which were lower in females than in males (p < 0.0001; **Fig. 6A**). While hepatic CAT protein was not impacted by birth weight nor sex, hepatic CAT activity was lower in male LBW (p < 0.05) (**Figs. 6A, 6B**). SOD activity was decreased in male LBW but increased in female LBW. (**Fig. 6C**). GSH activity was not significantly affected by birth weight or sex (**Fig. 6D**). While the concentration of GSSG was significantly reduced in male LBW males compared to male NBW (p <0.05; **Fig. 6E**), the ratio of GSH:GSSG was not significantly impacted by birth weight or sex (**Fig. 6F**).

**Fig. 6:**
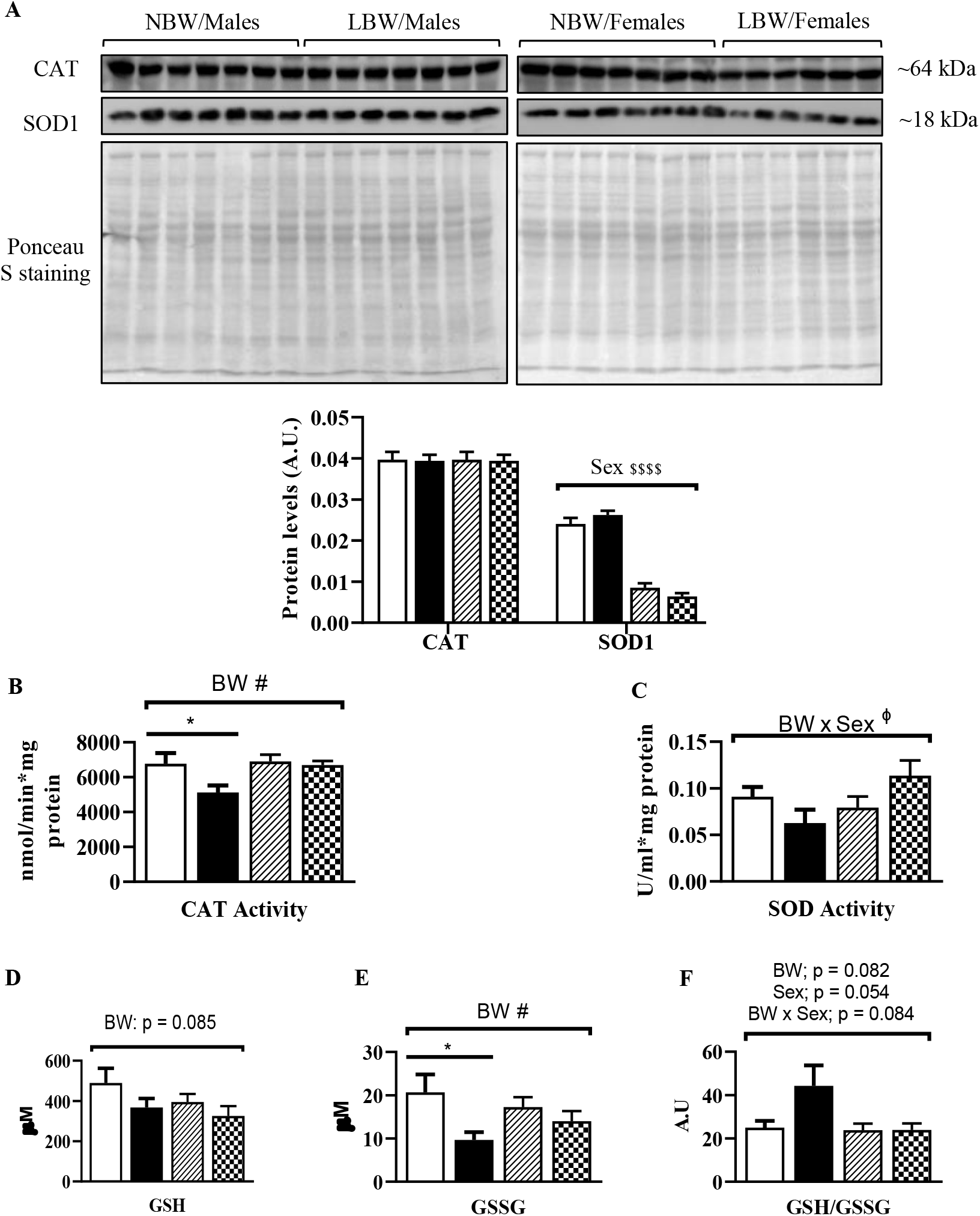
Immunoblotting and colorimetric assays of antioxidant system. Panel **A** shows representative blots of Catalase (CAT) and superoxide dismutase 1 (SOD1) proteins and ponceau as well as normalized densitometry values of targeted proteins. Panel **B** shows hepatic activity of catalase (CAT) and Panel **C** indicates hepatic activity of superoxide dismutase activity (SOD). Panels D, E and F show reduced glutathione (GSH), oxidized GSH (GSSG) and GSH/GSSG ratio, respectively. Data means ± SEM of 7 to 8 animals per birth weight/sex group. ^$$$$^p < 0.0001 for the main effect of sex; ^#^p < 0.05 or the indicated p-value for the main effect of birth weight (BW) and ^ϕ^p < 0.05 for the effect BW and sex interaction, using a two-way ANOVA. *p < 0.05 when comparing NBW/Males vs. LBW/Males by Bonferroni *post hoc* test.

## Discussion

An adverse environment during *in utero* life has been associated with gene reprogramming and modification of organ functions that can persist throughout the entire lifespan and are associated with an increased risk of developing the metabolic syndrome (47). Using a pre-clinical guinea pig animal model of UPI, we demonstrated that LBW is associated with increased hepatic cholesterol content and aberrant expression of cholesterol metabolism-related signalling pathway genes in liver. These changes occur in conjunction with markers of a compromised hepatic cholesterol elimination process and failing antioxidant system in young adults. Interestingly, these changes, at the current age studied, were sex-specific, only being observed in LBW males and not LBW females.

The impact of an adverse in utero environment on cholesterol metabolism has been mostly studied using mouse and rat models of IUGR arising from maternal protein undernutrition, *in utero* dietary restriction, prenatal hypoxia, or nicotine exposure (17–20). These different species and intrauterine insults collectively lead to elevated serum or hepatic cholesterol levels at weaning or in the adulthood in male offspring, similar to what was observed in this current study. The specific etiology of the IUGR insult is of critical importance in determining the adult metabolic outcomes and phenotype (22,23) and hence in the current study, UPI was investigated as it is the most common *in utero* insult associated with IUGR and LBW in developed world (48,49). Moreover, the guinea pig was utilized given its greater similarities with human pregnancy and outcomes including a relatively long gestation, haemomonochorial placenta, luteo-placental shift in hormone production, fetal development of metabolic tissues and precocial offspring (50). The reported outcomes here are similar to the observed increased in blood total cholesterol at 15 weeks of age observed in UPI-induced IUGR male rat offspring (51). Additionally, the observed increase in hepatic cholesterol without increased hepatic triglycerides in male LBW offspring is also in alignment with other species studies that report IUGR-induced alterations in the cholesterol metabolizing pathway, without changes in hepatic fatty acid metabolism (33). The current work highlights not only that the UPI environment is associated with programming of later life hepatic cholesterol dysfunction, but also validates the experimental *in utero* insults/stress situations used in other species. Collectively, this provides external validation for the central nature of this pathway and later life cholesterol metabolism dysregulation across species.

The present study also highlights that cholesterol metabolism and PPAR signalling pathways are impacted and associated genes are differentially expressed in LBW *versus* NBW male offspring, but not in females. Hepatic *Apoa1, Angplt4* and *Adipoq* genes, components of both cholesterol metabolism and PPAR pathways were up-regulated exclusively in LBW males. Apoa1 is the main apolipoprotein of HDL, playing a key role in regulating lipid transport and in the process of reverse cholesterol transport acting through promotion of the efflux of excess cholesterol from peripheral tissues and returning it to the liver for biliary excretion (52). Liver *ApoA-1* gene expression is up-regulated in liver disease states (53,54), and ApoA-1 protein among other liver-specific proteins in extracellular vesicles, has been suggested to potentially serve as a specific biomarker for hepatotoxicity in drug- and alcohol-mediated hepatic injury (55). Additionally, Angptl4 and Adipoq, like ApoA-1, collectively serve a number of important roles in cholesterol metabolism. Angptl4 is present in HDLs physically protecting HDLs from endothelial lipase hydrolysis (56), and upregulates cholesterol synthesis in liver secondary to inhibition of lipoprotein lipase- and hepatic lipase-dependent hepatic cholesterol uptake (57). In addition, Adipoq accelerates reverse cholesterol transport by increasing HDL assembly through enhanced ApoA-1 synthesis in the liver (58). Conversely, the decreased *Ldlr* gene expression and unaltered Ldlr protein in LBW males may reflect an unaltered LDL cholesterol uptake into these livers. In agreement with this assumption, it is well established that over-accumulation of cholesterol in the liver supresses *Ldlr* gene transcription and accelerates its mRNA decay (59). Finally, the intersection of cholesterol and PPAR pathways is of note given the relationship between PPARs in the regulation of pathways in bile acid and cholesterol homeostasis (60). Therefore, the current changes in gene expression collectively emphasize potential increased HDL assembly and/or reverse cholesterol transport, in conjunction with an unaltered LDL uptake in the LBW male livers, which elucidates the higher cholesterol content observed in these livers, though this remain to be more thoroughly investigated.

Although unaltered at transcriptional level, hepatic Cyp7a1 and Mtp was significantly reduced at the protein level in LBW males. The Cyp7a1 enzyme catalyzes the initial step in cholesterol catabolism and bile acid synthesis and decreased *Cyp7a1* gene expression and protein levels have been reported in conjunction with increased hepatic cholesterol in adult IUGR male rats from protein restricted mothers (17). Further, Cyp7a1 deficient mice are observed to have elevated hepatic and serum cholesterol and decreased total bile acids (61), supporting the concept that Cyp7A1 acts in a similar manner in guinea pigs and rats and that alterations in its level/activity impact hepatic cholesterol content. It is of further interest to note that the human *CYP7A1* mutation results in substantial cholesterol accumulation in the liver as well as decreased classic bile acid synthesis (62). Our current findings are in line with the concept that Cypa7a1 is critical in the control of hepatic cholesterol homeostasis and provide evidence that Cyp7a1-induced cholesterol catabolism appears sensitive to changes in the intrauterine environment, specifically those associated with UPI such as hypoxia and altered nutrient supply (17,19). The parallel decrease in Mtp protein is of great interest considering that Mtp catalyses the assembly of cholesterol, triglycerides, and apolipoprotein B to VLDL for their export outside of the liver (63). Therefore, this impairment in hepatic cholesterol catabolism/efflux-related proteins could reflect a lower elimination rate of hepatic cholesterol, programmed by an adverse *in utero* environment, which culminates in increased cholesterol content in LBW males born from pregnancies complicated by UPI.

Antioxidant enzyme defense and non-enzymatic antioxidant defense systems are critical in reducing oxidative stress and maintaining redox homeostasis within liver (64). SOD reduces the radical superoxide to form hydrogen peroxide and oxygen (65) and CAT catalytically decomposes hydrogen peroxide into water and oxygen (66). At the same time, glutathione peroxidase (GPx) also reduces hydrogen peroxide to water while converting GSH to GSSG (67). During oxidative stress there is decrease in levels of GSH and increase in levels of GSSG and thus GSH/GSSG ratio decreases (68). Liver oxidative injury and abnormal activities of hepatic antioxidant enzymes are reported in male neonates from IUGR pregnancies (69,70). In the current report, UPI-induced LBW male offspring displayed a significantly depressed CAT activity, and a reduced total SOD activity, a result not observed in the LBW female livers. The ratio of GSH to GSSG was however not reduced in the livers of LBW males. Collectively, these data suggest that whereas female offspring appear to have functional postnatal hepatic oxidative stress recovery mechanisms, the *in utero* defects in male offspring appear to persist, and this could be manifest with the altered hepatic cholesterol metabolic pathways observed. Indeed, in adult animal cholesterol feeding studies, elevated hepatic cholesterol is associated with oxidative stress (9,10). It is then possible that alterations in the functional postnatal hepatic oxidative stress of male LBW start at the level of SOD and CAT enzymes and may later extend to the glutathione redox couple GSH/GSSG system, as previously highlighted (68). This notion is further validated by the observed reduction in the expression of *inhba* and *Gstt2* genes, involved in hepatocyte regeneration and antioxidant system, respectively (71,72). Therefore, our current observations point to a lower antioxidant capacity in young adult LBW male offspring born from UPI pregnancies, which may promote the development of liver diseases in later life, especially when challenged with an elevated cholesterol environment.

In the current study, we observed lower body and liver weights at young adulthood in females than males, independent of the birth weight. Previous studies have also reported a sexual dimorphism in guinea pig with males consistently larger than females in skeletal measurements and body weight (73,74). It has been proposed that rapid and early growth in males leads to male-biased sexual dimorphism in these cases. In conjunction with these growth differences we report sexual dimorphism in hepatic Ldlr, Srebp2, Mtp, Acc and SOD1 proteins as well as liver cholesterol content, irrespective of birth weight, but also sex-specific programming of *Apoa1, Angplt4, and ldlr* genes and Cyp7a1 and Mtp proteins in male LBW offspring. Epidemiologic studies have demonstrated that LBW predisposes to adult onset hypercholesterolemia in men (26–28), and women (29) with sex difference by age groups in adults (24). In the current study, male offspring displayed higher hepatic free cholesterol and total cholesterol than females, despite higher protein abundance of Ldlr and Srebp2 and reduced Mtp and Acc proteins in female livers. We speculate that estrogens levels may be protective for cholesterol overaccumulation within livers of female guinea pigs, given that physiological levels of estrogen increased CYP7A1 activity along with small transient increases in BA production in hepatocytes (75). These results also indicate that such a protective mechanism may occur as a compensatory mechanism to higher protein level of Ldlr and Srebp2 in males and reduced Mtp and Acc proteins in females. It was interesting to note also the sex-specific difference observed in *Lcn2* and *Bcl6* in female LBW. These genes are involved in alleviating and promoting hepatic lipotoxicity, respectively (76,77), and their down-regulation in female LBW may underly a tight regulation of lipid catabolism in these offspring given their unaltered hepatic triglycerides and cholesterol contents.

The present study demonstrates that LBW occurring as a result of UPI results in increased hepatic cholesterol content, likely through compromised hepatic cholesterol elimination- in young male, but not female guinea pigs. The mechanisms underlying this potential programing effect and continued postnatal presentation of the defect are yet to be fully delineated. Certainly UPI is associated with hypoxia (78) and fetal hypoxia is associated with promoted oxygenated blood flow to the heart and reduced umbilical blood supply to fetal liver, an adaption which is understood to reprogram liver carbohydrate and lipid metabolism *in utero* (79–82) likely through altered epigenetic regulation of hepatic metabolism (17). Furthermore, the observed disrupted hepatic cholesterol metabolism may contribute to permanent alterations in hepatic oxidative stress defense system, ultimately leading to further hepatic damage and greater predisposition to liver diseases in UPI-induced LBW male offspring as they age.

## Author contributions

Ousseynou Sarr, and Timothy R.H. Regnault designed the experiments, analysed data and drafted the manuscript. Ousseynou Sarr, Katherine E. Mathers, Christina Vanderboor, Aditya Devgan, Lin Zhao and Timothy R.H. Regnault performed the experiments. All authors reviewed and edited the manuscript.

## Acknowledgments

the authors thank Brad Matushewski for his assistance with animal surgeries and Brian Sutherland for cholesterol measurements.

## Category of study

basic science.

## REFERENCES

1. Lin C, Lai C, Kao M, Wu L, Lo U, Lin L. Impact of cholesterol on disease progression. Biomedicine 2015;5:1–7.

2. Trapani L, Segatto M, Pallottini V, Trapani L, Segatto M, Pallottini V. Regulation and deregulation of cholesterol homeostasis: The liver as a metabolic “ power station World J Hepatol 2012;4:184–90.

3. Sharpe LJ, Brown AJ. Controlling Cholesterol Synthesis beyond 3-Hydroxy-3-methylglutaryl-CoA Reductase (HMGCR). J Biol Chem 2013;288:18707–15.

4. Feingold KR GC. Introduction to lipids and lipoproteins. In: Feingold KR, Anawalt B, Boyce A et al, editor. Diagnosis and Treatment of Diseases of Lipid and Lipoprotein Metabolism in Adults and Children. MDText.com, Inc., South Dartmouth (MA); 2000.

5. Zinkhan EK, Zalla JM, Carpenter JR, et al. Intrauterine growth restriction combined with a maternal high-fat diet increases hepatic cholesterol and low-density lipoprotein receptor activity in rats. Physiol Rep 2016;4:1–13.

6. Kosters A, Kunne C, Looije N, Patel SB, Elferink RPJO, Groen AK. The mechanism of ABCG5 / ABCG8 in biliary cholesterol secretion in mice 1. J Lipid Res 2006;47:1959–66.

7. Pandak WM, Schwarz C, Hylemon PB, et al. Effects of CYP7A1 overexpression on cholesterol and bile acid homeostasis. Am J Physiol Gastrointest Liver Physiol 2019;281:878–89.

8. Domínguez-pérez M, Nuño-lámbarri N, Clavijo-cornejo D, et al. Hepatocyte Growth Factor Reduces Free Cholesterol-Mediated Lipotoxicity in Primary Hepatocytes by Countering Oxidative Stress. Oxid Med Cell Longev 2016;2016:1–8.

9. Nuño-lámbarri N, Domínguez-pérez M, Baulies-domenech A, et al. Liver cholesterol overload aggravates obstructive cholestasis by inducing oxidative stress and premature death in mice. Oxid Med Cell Longev 2016;2016:1–13.

10. Pérez MD, Miranda RU, Bucio L, Carvajal SU, Marquardt JU, Quiroz LEG. Cholesterol burden in the liver induces mitochondrial dynamic changes and resistance to apoptosis. Cell Physiol 2019;234:7213–23.

11. López-islas A, Chagoya-hazas V, Pérez-aguilar B, et al. Cholesterol Enhances the Toxic Effect of Ethanol and Acetaldehyde in Primary Mouse Hepatocytes. Oxid Med Cell Longev 2016;2016:1–9.

12. López-Reyes A, Martínez-Flores K, Clavijo-Cornejo D, et al. Cholesterol overload in hepatocytes affects nicotinamide adenine dinucleotide phosphate oxidase (NADPH) activity abrogating hepatocyte growth factor (HGF) induced cellular protection. Gac Med Mex 2015;151:456–64.

13. Ioannou GN, Haigh WG, Thorning D, Savard C. Hepatic cholesterol crystals and crown-like structures distinguish NASH from simple steatosis. J Lipid Res 2013;54:1326–1334.

14. Ioannou G, Landis C, Jin G, et al. Cholesterol crystals in hepatocyte lipid droplets are strongly associated with human nonalcoholic steatohepatitis. Hepatol Commun 2019;3:776–91.

15. Paththinige CS, Sirisena ND, Dissanayake VHW. Genetic determinants of inherited susceptibility to hypercholesterolemia - a comprehensive literature review. Lipids Health Dis 2017;16:1–22.

16. Bellanti F, Villani R, Tamborra R, et al. Redox Biology Synergistic interaction of fatty acids and oxysterols impairs mitochondrial function and limits liver adaptation during na fl d progression. Redox Biol 2018;15:86–96.

17. Sohi G, Marchand K, Revesz A, Arany E, Hardy DB. Maternal protein restriction elevates cholesterol in adult rat offspring due to repressive changes in histone modifications at the cholesterol 7alpha-hydroxylase promoter. Mol Endocrinol 2011;25:785–98.

18. Zhou J, Zhu C, Luo H, et al. Two intrauterine programming mechanisms of adult hypercholesterolemia induced by prenatal nicotine exposure in male offspring rats. FASEB J 2019;33:1110–23.

19. Rueda-clausen CF, Dolinsky VW, Morton JS, Proctor SD, Dyck JRB, Davidge ST. Hypoxia-induced intrauterine growth restriction increases the susceptibility of rats to high-fat diet-induced metabolic syndrome. Diabetes 2011;60:507–16.

20. Choi GY, Tosh DN, Garg A, Mansano R, Ross MG, Desai M. Gender-specific programmed hepatic lipid dysregulation in intrauterine growth-restricted offspring. Am J Obs Gynecol 2007;196:1–7.

21. Lecoutre S, Montel V, Vallez E, et al. Transcription profiling in the liver of undernourished male rat offspring reveals altered lipid metabolism pathways and predisposition to hepatic steatosis. Am J Physiol Endocrinol Metab 2019;317:E1094–107.

22. Nüsken KD, Schneider H, Plank C, et al. Fetal programming of gene expression in growth-restricted rats depends on the cause of low birth weight. Endocrinology 2011;152:1327–35.

23. Nüsken E, Fink G, Lechner F, et al. Altered molecular signatures during kidney development after intrauterine growth restriction of different origins. J Mol Med 2020;98:395–407.

24. Chen L, Chen S, Liang L, et al. Relationship between birth weight and total cholesterol concentration in adulthood: A meta-analysis. J Chinese Med Assoc 2017;80:44–9.

25. Barker DJP, Martyn CN, Osmond C, Hales CN, Fall CHD. Growth in utero and serum cholesterol concentrations in adult life. BMJ 1993;307:1524–1527.

26. Ziegler B, Johnsen SP, Thulstrup AM, Engberg M, Lauritzen T, Sørensen HT. Inverse association between birth weight, birth length and serum total cholesterol in adulthood. Scand Cardiovasc J 2000;34:584–8.

27. Cooper, Rachel1. Cooper R PCS differences in the associations between birthweight and lipid levels in middle-age: F from the 1958 B birth cohort. A 2008;200:141–9.,

28. Power C. Sex differences in the associations between birthweight and lipid levels in middle-age: Findings from the 1958 British birth cohort. Atherosclerosis 2008;200:141–9.

28. Davies AA, Smith GD, Ben-Shlomo Y, Litchfield P. Low birth weight is associated with higher adult total cholesterol concentration in men: Findings from an occupational cohort of 25 843 employees. Circulation 2004;110:1258–62.

29. Miura K, Nakagawa H, Tabata M, Morikawa Y, Nishijo M KS. Birth Weight, Childhood Growth, and Cardiovascular Disease Risk Factors in Japanese Aged 20 Years. Am J Epidemiol 2001;153:783–9.

30. Henriksen T, Clausen T. The fetal origins hypothesis: Placental insufficiency and inheritance versus maternal malnutrition in well-nourished populations. Acta Obstet Gynecol Scand 2002;81:112–4.

31. Ciurea EL, Berceanu C, Voicu NL, Pirnoiu D, Berceanu S, Stepan AE. Morphological Survey of Placenta in Trombophilia Related Hypoperfusion of Maternal-Fetal Blood Flow. Curr Heal Sci J 2018;44:85–91.

32. Nüsken K, Dötsch J, Rauh M, Rascher W, Schneider H. Uteroplacental insufficiency after bilateral uterine artery ligation in the rat: impact on postnatal glucose and lipid metabolism and evidence for metabolic programming of the offspring by sham operation. Endocrinology 2008;149:1056–63.

33. Zinkhan EK, Chin JR, Zalla JM, et al. Combination of intrauterine growth restriction and a high-fat diet impairs cholesterol elimination in rats. Pediatr Res 2014;76:432–40.

34. Miranda J, Simões R V., Paules C, et al. Metabolic profiling and targeted lipidomics reveals a disturbed lipid profile in mothers and fetuses with intrauterine growth restriction. Sci Rep 2018;8:1–14.

35. Turner AJ, Trudinger BJ. A modification of the uterine artery restriction technique in the guinea pig fetus produces asymmetrical ultrasound growth. Placenta 2009;30:236–40.

36. Fernandez ML, Volek JS. Guinea pigs: A suitable animal model to study lipoprotein metabolism, atherosclerosis and inflammation. Nutr Metab 2006;3:1–6.

37. Grundy D. Principles and standards for reporting animal experiments in The Journal of Physiology and Experimental Physiology. J Physiol 2015;593:2547–9.

38. Kilkenny C, Browne WJ, Cuthill IC, Emerson M, Altman DG. Improving Bioscience Research Reporting: The ARRIVE Guidelines for Reporting Animal Research. PLoS Biol 2010;8:e1000412.

39. Sarr O, Thompson JA, Zhao L, Lee TY, Regnault TRH. Low birth weight male guinea pig offspring display increased visceral adiposity in early adulthood. PLoS One 2014;9:1–13.

40. Thompson JA, Gros R, Richardson BS, Piorkowska K, Regnault TRH. Central stiffening in adulthood linked to aberrant aortic remodeling under suboptimal intrauterine conditions. AJP Regul. Integr. Comp. Physiol. 2011;301:R1731–7.

41. Greulich S, Herzfeld D, Wiza D, et al. Secretory products of guinea pig epicardial fat induce insulin resistance and impair primary adult rat cardiomyocyte function. J Cell Mol Med 2011;15:2399–410.

42. Samsoondar JP, Burke AC, Sutherland BG, et al. Prevention of diet-induced metabolic dysregulation, inflammation, and atherosclerosis in Ldlr-/-mice by treatment with the ATP-citrate lyase inhibitor bempedoic acid. Arter Thromb Vasc Biol 2017;37:647–56.

43. Folch J, Lees M, Sloane Stanley GH. A simple method for the isolation and purification of total lipides from animal tissues. J Biol Chem 1957;226:497–509.

44. Sarr O, Mathers KE, Zhao L, et al. Western diet consumption through early life induces microvesicular hepatic steatosis in association with an altered metabolome in low birth weight Guinea pigs. J Nutr Biochem 2019;67:219–33.

45. Lojpur T, Easton Z, Raez-Villanueva S, Laviolette S, Holloway AC, Hardy DB. Δ9-Tetrahydrocannabinol leads to endoplasmic reticulum stress and mitochondrial dysfunction in human BeWo trophoblasts. Reprod Toxicol 2019;87:21–31.

46. Sander H, Wallace S, Plouse R, Tiwari S, Gomes A V. Ponceau S waste: Ponceau S staining for total protein normalization. Anal Biochem 2019;575:44–53.

47. Gluckman P, Hanson M, Phil D, Cooper C, Thornburg K. Effect of In Utero and Early-Life Conditions on Adult Health and Disease. N Engl J Med 2014;359:61–73.

48. Brown LD, Hay WW. Impact of placental insufficiency on fetal skeletal muscle growth. Mol Cell Endocrinol 2017;435:69–77.

49. Kinzler WL, Vintzileos AM. Fetal growth restriction: a modern approach. Curr Opin Obs Gynecol 2008;2:125–31.

50. Morrison JL, Botting KJ, Darby JRT, et al. Guinea pig models for translation of the developmental origins of health and disease hypothesis into the clinic. J Physiol 2018;23:5535–69.

51. Nüsken KD, Dötsch J, Rauh M, Rascher W, Schneider H. Uteroplacental insufficiency after bilateral uterine artery ligation in the rat: Impact on postnatal glucose and lipid metabolism and evidence for metabolic programming of the offspring by sham operation. Endocrinology 2008;149:1056–63.

52. Lewis GF, Rader DJ. New insights into the regulation of HDL metabolism and reverse cholesterol transport. Circ Res 2005;96:1221–32.

53. Panduro A, Castrillb L, Gonzalez L, Shafritz DA. Regulation of hepatic and non-hepatic apolipoprotein gene expression during liver regeneration A-I and E. Biochim Biophys Acta 1993;1167:37–42.

54. Hilippe P, Idaud DO V, Idaud MI V, Edossa PIB, Ale V. Quantification of apolipoprotein A-I and B messenger RNA in heavy drinkers according to liver disease. Hepatology 1996;23:44–51.

55. Cho Y, Im E, Moon P, Mezey E, Song B, Baek M. Increased liver-specific proteins in circulating extracellular vesicles as potential biomarkers for drug-and alcohol-induced liver injury. PLoS One 2017;12:1–23.

56. Yang L, Yu C, Wang X, et al. Angiopoietin-like protein 4 is a high-density lipoprotein (HDL) component for HDL metabolism and function in nondiabetic participants and type-2 diabetic patients. J Am Hear Assoc 2017;6:1–15.

57. Dijk SJ Van, Dijk KW Van, Kema IP, et al. Angptl4 upregulates cholesterol synthesis in liver via inhibition of LPL-and HL-dependent hepatic cholesterol uptake. Arter Thromb Vasc Biol 2007;27:2420–7.

58. Matsuura F, Oku H, Koseki M, et al. Adiponectin accelerates reverse cholesterol transport by increasing high density lipoprotein assembly in the liver. Biochem Biophys Res Commun 2007;358:1091–5.

59. Singh AB, Fung C, Kan K, Shende V, Dong B, Liu J. A novel posttranscriptional mechanism for dietary cholesterol-mediated suppression of liver LDL receptor expression. J Lipid Res 2014;55:1397–407.

60. Li, Tiangang, Chiang JYL. Regulation of bile acid and cholesterol metabolism by PPARs. PPAR Res 2009;2009.

61. Erickson SK, Lear SR, Deane S, et al. Hypercholesterolemia and changes in lipid and bile acid metabolism in male and female cyp7A1-deficient mice. J Lipid Res 2003;44:1001–9.

62. Pullinger CR, Eng C, Salen G, et al. Human cholesterol 7 α - hydroxylase (CYP7A1) deficiency has a hypercholesterolemic phenotype. J Clin Invest 2002;110:109–117.

63. Gordon DA, Jamil H. Progress towards understanding the role of microsomal triglyceride transfer protein in apolipoprotein-B lipoprotein assembly. Biochim Biophys Acta 2000;1486:72–83.

64. Li S, Tan H, Wang N, et al. The role of oxidative stress and antioxidants in liver diseases. Int J Mol Sci 2015;16:26087–124.

65. Buettner R, Parhofer KG, Woenckhaus M, et al. Defining high-fat-diet rat models: metabolic and molecular effects of different fat types. J Mol Endocrinol 2006;36:485–501.

66. Patlevi P, Svorc P. Reactive oxygen species and antioxidant defense in human gastrointestinal diseases. Integr Med Res 2016;5:250–8.

67. Aquilano K, Baldelli S, Ciriolo MR. Glutathione: New roles in redox signalling for an old antioxidant. Front Pharmacol 2014;5 AUG:1–12.

68. Raab EL, Vuguin PM, Stoffers DA, Simmons RA. Neonatal exendin-4 treatment reduces oxidative stress and prevents hepatic insulin resistance in intrauterine growth-retarded rats. Am J Physiol - Regul Integr Comp Physiol 2009;297.

69. Hu L, Peng X, Qin L, et al. Advances dietary nucleotides supplementation during the suckling period improves the antioxidative ability of neonates with intrauterine growth retardation when using a pig model. RSC Adv 2018;8:16152–60.

70. Liu J, Yao Y, Yu B, Mao X, Huang Z, Chen D. Effect of folic acid supplementation on hepatic antioxidant function and mitochondrial-related gene expression in weanling intrauterine growth retarded piglets. Livest Sci 2012;146:123–32.

71. Kreidl E, Öztürk D, Metzner T, et al. Activins and follistatins: Emerging roles in liver physiology and cancer. World J Hepatol 2009;1:17–27.

72. Xu S, Hou D, Liu J, Ji L. Communications in Free Radical Research Age-associated changes in GSH S-transferase gene / proteins in livers of rats. Redox Rep 2018;23:213–8.

73. Farmer MA, German RZ. Sexual dimorphism in the craniofacial growth of the guinea Pig (Cavia porcellus). J Morphol 2004;259:172–81.

74. Oliver A, Norbert S. Diversity of social and mating systems in cavies: a review. J Mammal 2011;92:39–53.

75. Phelps T, Snyder E, Rodriguez E, Child H, Harvey P. The influence of biological sex and sex hormones on bile acid synthesis and cholesterol homeostasis. Biol Sex Differ 2019;10:1–12.

76. LaPensee CR, Lin G, Dent AL, Schwartz J. Deficiency of the transcriptional repressor B cell lymphoma 6 (Bcl6) is accompanied by dysregulated lipid metabolism. PLoS One 2014;9.

77. Asimakopoulou A, Weiskirchen R. Lipocalin 2 in the pathogenesis of fatty liver disease and nonalcoholic steatohepatitis. Clin Lipidol 2015;10:47–67.

78. Hutter D, Kingdom J, Jaeggi E. Causes and mechanisms of intrauterine hypoxia and its impact on the fetal cardiovascular system: a review. Int J Pediatr 2010;2010:401323.

79. Reuss ML RA. Distribution and recirculation of umbilical and systemic venous blood flow in fetal lambs during hypoxia. J Dev Physiol 1980;2:71–81.

80. Ring JA, Ghabrial H, Ching MS, Smallwood RA, Morgan DJ. Fetal hepatic drug elimination. Pharmacol Ther 1999;84:429–45.

81. Haugen G, Kiserud T, Godfrey K, Crozier S, Hanson M. Portal and umbilical venous blood supply to the liver in the human fetus near term. Ultrasound Obs Gynecol 2004; 2004;24:599–605.

82. Tchirikov M, Hecher K. Ductus venosus shunting in the fetal venous circulation: regulatory mechanisms, diagnostic methods and medical importance. Ultrasound Obs Gynecol 2006;27:452–61.

