## Supplementary Tables for "Sex-specific Alterations in Hepatic Cholesterol Metabolism in Young Uteroplacental Insufficiency-induced Low Birth Weight Adult Guinea Pig Offspring"

**Supplementary Table1:** Primer sequences used to measure guinea pig hepatic mRNA expressions by RTq-PCR

| Gene | Accession number * | Annealing temperature | Forward/reverse primer |
|---|---|---|---|
| *Apoa1* | XM_013155451.1 | 59 ºC | Forward: 5'-CTGAGGCTTTCGGATAACTG-3'<br>Reverse: 5'-TTCTTTTTCCAAGGTGTCCC-3' |
| *Angplt4* | XM_003461395.3 | 58 ºC | Forward: 5'-GAAGCAGCACTTGAGAATCA-3'<br>Reverse: 5'-AAACCGGGTTATCTTGAGGA-3' |
| *Anxa1* | XM_003472219.3 | 58 ºC | Forward: 5'-CTGGAAAGGTGTGGATGAG-3'<br>Reverse: 5'- TCAGTTCCAAGTCCCTTCAT-3' |
| *Ldlr* | XM_013149927 | 57 ºC | Forward: 5'-CATCTACTCGCTTGTGACAG-3'<br>Reverse: 5'-CTCATCCTCCAAGATGGTCT-3' |
| *Gstt2* | XM_003477956 | 58 ºC | Forward: 5'-GATGCAGCCAGTGATTTTTG-3'<br>Reverse: 5'-GAAGCAGGAAACTTGGGTAG-3' |
| *Inhba* | XM_013152606 | 54 ºC | Forward: 5'-TGTAGACTAGCACAACCCAA-3'<br>Reverse: 5'-AGCAGTATGAGCATGTAGGT-3' |
| *Lbp* | XM_003467651 | 54 ºC | Forward: 5'-AGGAAAATGTTACAGCGCCT-3'<br>Reverse: 5'-TCCGGAACATTTGTGCTTTC-3' |
| *Lnc2* | XM_005004004 | 58 ºC | Forward: 5'-ACACCCCCTTCTTTCTGTTA-3'<br>Reverse: 5'-AAGAGGGTTCCATCAGTAGG-3' |
| *Bcl6* | XM_013160235 | 60 ºC | Forward: 5'-CGAACTCCCTGACTCTATGA-3'<br>Reverse: 5'-AAGAACTAAACCGAAAGCCC-3' |
| *Gapdh* | NM_001172951.1 | 54 ºC | Forward: 5'- GCTCGTTTCTTGGTATGACA -3'<br>Reverse: 5'- CTAGTCTCCATGGTCTCACT -3' |
| *B-actin* | NM_001172909.1 | 62 ºC | Forward: 5'- AAGAGATGTGGCCTCAAAGC-3'<br>Reverse: 5'- CAGGAACAGGCCGTAGAGTG-3' |

**Supplementary Table 2**: Differentially expressed genes in male LBW compared to male NBW guinea pigs in Liver

| Transcript ID | Gene Symbol | Fold Change | P-value | Description |
|---|---|---|---|---|
| 20160052 | *Cxcl17* | -45.34 | 0.0009 | Chemokine (C-X-C motif) ligand 17 |
| 20051023 | *LOC106027169* | -4.29 | 0.0423 | Chromosome unknown open reading frame, human C19orf80 |
| 20017669 | *39d1* | -3.04 | 0.0354 | Enterophilin-1 |
| 20079533 | *Inhba* | -2.82 | 0.0362 | Inhibin, beta A |
| 20166493 | *Serpine2* | -2.67 | 0.0056 | Serpin peptidase inhibitor, clade E (nexin, plasminogen activator inhibitor type 1), member 2 |
| 20171290 | *Inhbe* | -2.51 | 0.0047 | Inhibin, beta E |
| 19969683 | *Nkain3* | -2.48 | 7.39E-06 | Na+/K+ transporting ATPase interacting 3 |
| 20017679 | *Btnl1* | -2.46 | 0.0234 | Butyrophilin-like protein 1 |
| 20095756 | *Gstt2* | -2.44 | 0.0412 | Glutathione S-transferase theta-2 |
| 20051006 | *Ldlr* | -2.39 | 0.0206 | Low density lipoprotein receptor |
| 20081028 | *SNORA77* | -2.37 | 0.0471 | Small nucleolar RNA SNORA77 |
| 20170100 | *Slc17a8* | -2.37 | 0.0066 | Solute carrier family 17 (vesicular glutamate transporter), member 8 |
| 20172505 | *Krtap12-1* | -2.34 | 0.0165 | Keratin-associated protein 12-3-like |
| 20087290 | *Mid1ip1* | -2.32 | 0.0062 | MID1 interacting protein 1 |
| 20142060 | *ENSCPOT00000010959* | -2.24 | 0.0148 | Novel protein coding gene |
| 20004001 | *ENSCPOT00000026779* | -2.24 | 0.0157 | Novel protein coding gene |
| 20079249 | *snoU13* | -2.21 | 0.0023 | Small nucleolar RNA U13-201 |
| 20116118 | *ENSCPOT00000002339* | -2.09 | 0.0444 | Novel protein coding gene |
| 20107579 | *5S_rRNA* | -2.08 | 0.0472 | 5S ribosomal RNA-201 |
| 19982082 | *5S_rRNA* | -2.06 | 0.003 | 5S ribosomal RNA |
| 20111111 | *Zbtb3* | -2.04 | 0.0453 | Zinc finger and BTB domain containing 3 |
| 20135848 | *LOC100713908* | -2.04 | 0.036 | Olfactory receptor 6C74-like |
| 20005858 | *nell1* | 2.01 | 0.0416 | Protein kinase C-binding protein NELL1 |
| 20010003 | *Nedd4l* | 2.02 | 0.0479 | Neural precursor cell expressed, developmentally down-regulated 4-like, E3 ubiquitin protein ligase |
| 20053812 | *Tubb3* | 2.03 | 0.0153 | Tubulin, beta 3 class III |
| 20153038 | *Cog1* | 2.03 | 0.0006 | Conserved oligomeric Golgi complex subunit 1-like |
| 20056640 | *ENSCPOT00000019214* | 2.04 | 0.0083 | Novel miRNA gene |
| 20078511 | *PilrbL* | 2.07 | 0.0401 | Paired immunoglobulin-like type 2 receptor beta |
| 20162740 | *5S_rRNA* | 2.08 | 0.0365 | 5S ribosomal RNA |
| 19989086 | *SNORA2* | 2.09 | 0.0055 | Small nucleolar RNA SNORA2/SNORA34 family |
| 19974163 | *ENSCPOT00000005958* | 2.13 | 0.0096 | Novel protein coding gene |
| 20176038 | *Txnp1* | 2.13 | 0.0093 | Thioredoxin pseudogene |
| 20117629 | *Slc45a4* | 2.16 | 0.0029 | Solute carrier family 45 member 4 |
| 20120669 | *Ica1* | 2.17 | 8.08E-05 | Islet cell autoantigen 1 |
| 20046344 | *Anxa1* | 2.18 | 0.0147 | Annexin A1 |
| 20137828 | *Ace* | 2.21 | 0.0002 | Angiotensin I converting enzyme |
| 20096528 | _ | 2.26 | 0.0018 | _ |
| 20050642 | *Mt2a* | 2.3 | 0.0252 | Metallothionein-2 |
| 20046045 | _ | 2.33 | 0.0466 | _ |
| 20140096 | *SNORA23* | 2.35 | 0.0101 | Small nucleolar RNA SNORA23 |
| 20170753 | *snoU13* | 2.53 | 0.0062 | Small nucleolar RNA U13 |
| 20114709 | *Ugt2a1* | 2.67 | 0.0188 | UDP glucuronosyltransferase 2 family, polypeptide A1, complex locus |
| 20143911 | *SNORA63* | 2.82 | 0.007 | Small nucleolar RNA SNORA63 |
| 20036817 | *Angptl4* | 2.84 | 0.0111 | Angiopoietin-like 4 |
| 20143915 | *Adipoq* | 2.9 | 0.0493 | Adiponectin, C1Q and collagen domain containing |
| 20034318 | *Apoa1* | 2.92 | 0.0366 | Apolipoprotein A-I |
| 20054323 | *SNORA49* | 3.07 | 0.0174 | Small nucleolar RNA SNORA49 |
| 19966934 | *Hey1* | 3.21 | 4.42E-06 | Hes-related family bHLH transcription factor with YRPW motif 1 |
| 20178145 | *LOC106024976* | 3.26 | 0.048 | Metallothionein-2-like |
| 19973535 | *Nrep* | 3.27 | 0.0124 | Neuronal regeneration related protein |
| 20010918 | *SNORA2* | 3.32 | 0.0241 | Small nucleolar RNA SNORA2/SNORA34 family |

Fold changes preceded by negative signs indicate downregulated genes whereas positive fold changes represent upregulated genes in LBW *versus* NBW livers.

**Supplementary Table 3:** Differentially expressed genes in female LBW compared to female NBW guinea pigs in Liver

| Transcript | Gene Symbol | Fold Change | P-value | Description |
|---|---|---|---|---|
| 20141168 | *LOC100718750* | -5.62 | 0.0005 | serum amyloid A-3 protein-like |
| 20038746 | *LOC100725801* | -5.51 | 0.017 | 3 beta-hydroxysteroid dehydrogenase/Delta 5-->4-isomerase-like |
| 20042590 | *LOC100713524* | -3.06 | 0.0023 | alpha-1-acid glycoprotein 1-like |
| 20122545 | *SNORA2* | -2.69 | 0.0002 | Small nucleolar RNA SNORA2/SNORA34 family |
| 20119786 | *Lrg1* | -2.69 | 0.0033 | leucine-rich alpha-2-glycoprotein 1 |
| 20066246 | *LOC100724477* | -2.68 | 0.0083 | neutrophil gelatinase-associated lipocalin |
| 20078633 | *Cyp3a17* | -2.62 | 0.0033 | cytochrome P450 3A17 |
| 20145946 | *Bcl6* | -2.55 | 0.0135 | B-cell CLL/lymphoma 6 |
| 20095774 | *LOC100717598* | -2.52 | 0.0332 | glutathione S-transferase theta-1-like |
| 20093646 | *Adtrp* | -2.51 | 0.0142 | androgen-dependent TFPI-regulating protein |
| 20056757 | _ | -2.49 | 0.0186 | _ |
| 20170852 | *LOC100732231* | -2.45 | 0.03 | lysozyme C |
| 20010918 | *SNORA2* | -2.43 | 0.0206 | Small nucleolar RNA SNORA2/SNORA34 family |
| 20023554 | *Gadd45g* | -2.42 | 0.0201 | growth arrest and DNA-damage-inducible, gamma |
| 20018367 | *SNORD116* | -2.4 | 0.018 | small nucleolar RNA SNORD116 |
| 20146690 | *LOC106026670* | -2.35 | 0.024 | chromosome unknown open reading frame, human C19orf84 |
| 20044954 | *U6* | -2.31 | 0.0475 | U6 spliceosomal RNA |
| 20112268 | *Lbp* | -2.28 | 0.0257 | lipopolysaccharide binding protein |
| 20010916 | *SNORA2-201* | -2.19 | 0.0106 | Small nucleolar RNA SNORA2/SNORA34 family |
| 20141166 | *SNORA42* | -2.17 | 0.0012 | small nucleolar RNA SNORA42/SNORA80 |
| 20176038 | *LOC100713255* | -2.09 | 0.0123 | thioredoxin pseudogene |
| 20008495 | _ | -2.09 | 0.0104 | _ |
| 20043225 | *Scarna11* | -2.02 | 0.0024 | Small Cajal body specific RNA 11 |
| 20103473 | *Taf1d* | 2.08 | 0.0165 | TATA box binding protein (TBP)-associated factor, RNA polymerase I, D, 41kDa |
| 20000069 | *Serpina6* | 2.09 | 0.0135 | serpin peptidase inhibitor, clade A (alpha-1 antiproteinase, antitrypsin), member 6 |
| 20140350 | *Insc* | 2.34 | 0.0486 | inscuteable homolog (Drosophila) |
| 20156307 | *SNORA2* | 2.46 | 0.0268 | Small nucleolar RNA SNORA2/SNORA34 family |
| 20133878 | *Linc00675* | 2.65 | 0.0293 | long intergenic non-protein coding RNA 675 |
| 20135029 | *U6atac* | 2.82 | 0.0114 | U6atac minor spliceosomal RNA |
| 20178753 | *SNORA2-201* | 3.17 | 0.0055 | Small nucleolar RNA SNORA2/SNORA34 family |
| 20086987 | *Agpat9* | 3.68 | 0.0317 | 1-acylglycerol-3-phosphate O-acyltransferase 9 |

Fold changes preceded by negative signs indicate downregulated genes whereas positive fold changes represent upregulated genes in LBW *versus* NBW livers.
